# SWEET family transporters act as water conducting carrier proteins in plants

**DOI:** 10.1101/2024.06.23.600272

**Authors:** Balaji Selvam, Arnav Paul, Ya-Chi Yu, Li-Qing Chen, Diwakar Shukla

## Abstract

Dedicated water channels are involved in the facilitated diffusion of water molecules across the cell membrane in plants. Transporter proteins are also known to transport water molecules along with substrates, however the molecular mechanism of water permeation is not well understood in plant transporters. Here, we show plant sugar transporters from the SWEET (**S**ugar **W**ill **E**ventually be **E**xported **T**ransporter) family act as water-conducting carrier proteins via a variety of passive and active mechanisms that allow diffusion of water molecules from one side of the membrane to the other. This study provides a molecular perspective on how plant membrane transporters act as water carrier proteins, a topic that has not been extensively explored in literature. Water permeation in membrane transporters could occur via four distinct mechanisms which form our hypothesis for water transport in SWEETs. These hypothesis are tested using molecular dynamics simulations of the outward-facing, occluded, and inward-facing state of AtSWEET1 to identify the water permeation pathways and the flux associated with them. The hydrophobic gates at the center of the transport tunnel act as a barrier that restricts water permeation. We have performed *in silico* single and double mutations of the hydrophobic gate residues to examine the changes in the water conductivity. Surprisingly, the double mutant allows the water permeation to the intracellular half of the membrane and forms a continuous water channel. These computational results are validated by experimentally examining the transport of hydrogen peroxide molecules by the AtSWEET family of transporters. We have also shown that the transport of hydrogen peroxide follows the similar mechanism as water transport in AtSWEET1. Finally, we conclude that similar water-conduction states are also present in other SWEET transporters due to the high sequence and structure conservation exhibited by this transporter family.

## Introduction

Water is crucial for all cellular chemical reactions and physiological processes, determining plant growth and development. Dedicated water channels such as Aquaporins, facilitate water movement across cell membranes and play a fundamental role in various physiological processes.^1^ These water channels have been identified in mammals, bacteria, yeast and amphibians. In plants, plasma membrane localized and tonoplast localized water channels have been characterized.^2–4^ In addition to channels, membrane transporters have been shown to facilitate water permeation but such observations have not been reported for plants.^5–7^ Recent findings suggest that substrate transport in secondary active transporters leads to water transport in the direction of substrate transport. The water conducting states are well-documented in sodium-galactose cotransporter (SGLT1),^8–11^ glucose transporter (GLUT1),^12^ potassium-chloride cotransporter (KCC), ^13^ lactate transporter^14^ and LeuT-fold transporters.^15^ Transporters are also shown to form transient water conducting channels.^16^ The molecular mechanism of water permeation and the associated pathways in membrane transporters are not as well characterized as in channels because membrane transporters cycle through multiple structural conformational states necessary for transport whereas channels demonstrate minimal conformational variability.

In this study, we demonstrate that a plant sugar transporter family, SWEET (**S**ugar **W**ill **E**ventually be **E**xported **T**ransporter)^17–19^ also acts as a water-conducting carrier proteins in plants. Sugar transporters play an important role in translocating sugars across different cells or different cellular compartments. In our previous studies, the functionally important intermediate states of SWEET^20–23^ and SemiSWEET^24,25^ transporters as well as the complete cycle of sugar transport has been characterized using large-scale Molecular Dynamics (MD) simulations and Markov State Modeling of the transporter dynamics. Markov state modeling of membrane transporters quantifies the thermodynamics and kinetics associated with the transports process and aids in the investigation of structure-function relationship of the complex membrane proteins.^26–31^

Previous studies have suggested that water gets transported along with the substrates in sugar transporters such as SGLT1 and GLUT1 via distinct transport mechanisms.^8,32–34^

Experimental studies suggest that SGLT transports *∼*175 molecules molecules per ion or other substrate to regulate the osmotic concentration of the cell membrane. ^35^ In this work, we propose four hypotheses for water transport mechanisms in SWEETs. **Hypothesis 1:** Substrate water cotransport model, where water molecules get cotransported along with the substrate (Fig. 1A). **Hypothesis 2:** Apo-transporter water permeation model, the apo transporter completes an alternating access cycle without substrate involved and only carries water across the membrane (Fig. 1B). **Hypothesis 3:** Water escape model, the water translocation pore is open wide enough to allow water to escape to the intracellular side (Fig. 1C). **Hypothesis 4:** Water channel-like model, a continuous water channel that allows passive water transport (Fig. 1D).

**Figure 1:**
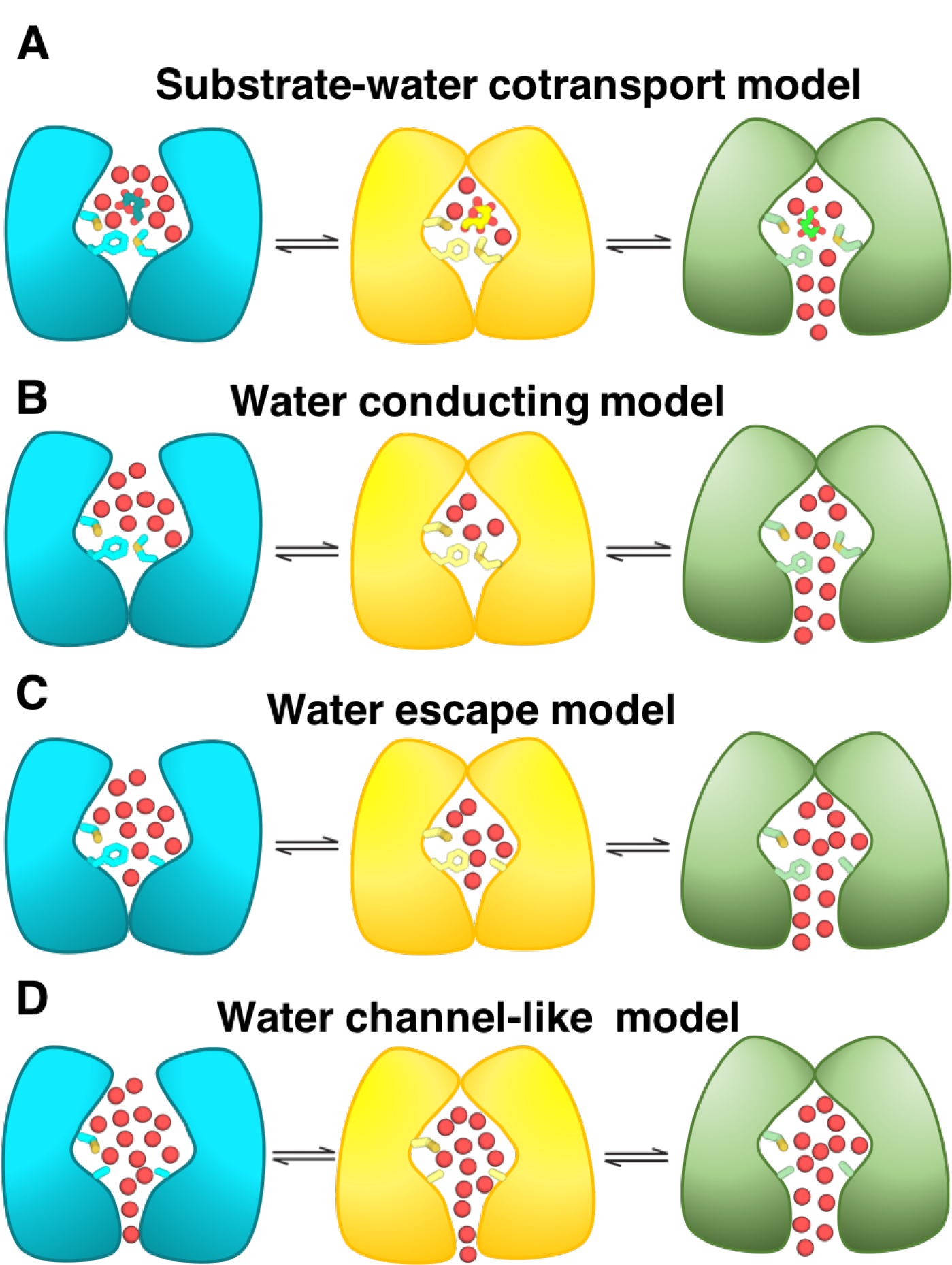
Water permeation models for membrane transporters. A) The substrate coupled water permeation of transporters. B) Water conducting states of apo transporter. C) Wide translocation pore allows water to escape to the intracellular side and D) Translocation pore wide enough for channel like formation for water transport.

To test these hypotheses, the functionally important conformational states in the transport cycle i.e. Outward facing (OF), Occluded (OC) and Inward facing (IF) conformations of AtSWEET1 were used to perform MD simulation for probing the water transport mechanism and its flux. Our results show that holo AtSWEET1 follows the substrate-water cotransport model where water molecules surrounding the sugars in the binding cavity are co-transported with the substrate. The apo AtSWEET1 follows the water conducting model where the water is transported due to the water filled binding site in the OC state. We also show that the other water permeation mechanisms are feasible when mutation are introduced in the hydrophobic gating residues. These gating residues are known to restrict sugar entry to the cytoplasmic portion of the protein channel within the OF and OC states.^20,21^ In particular, the double mutation in these gates leads to the formation of a continuous water channel and allows for exchange of water from the extracellular to intracellular side between the OF and OC states. These results have been validated experimentally using yeast growth assays which show that SWEETs transport (H_2_O_2_), which served as an analog of water as they posses similar physical properties. We also provide a molecular basis of SWEETs facilitating passive diffusion of hydrogen peroxide (H_2_O_2_), suggesting a crucial role in plant signaling, growth and development in addition to sugar transport. Finally, this study establishes a new transporter family in plants as water-conducting channels and provides avenues to better understand the plant membrane transport processes.

## Methods

### Simulation Details

The outward-facing (OF), occluded (OC) and inward-facing (IF) states of OsSWEET2b structures were used as structural templates for modeling the AtSWEET1 structures (Fig. 2). Alphafold2^36^ could only predict one conformational state of AtSWEET1 and several regions in the predicted structure had low pLDDT score, so the output structure from AlphaFold2 was not used for simulations in this study. The homology models were built using MODELLER.^37^ The MD system was built using CHARMM-GUI^38^ and converted to amber format files using Tleap program in AMBER.^39^ The default force field amberff14SB was used for MD simulation.^40^ The OF, OC and IF states were embedded in a phospholipid bilayer and solvated using TIP3P water molecules. NaCl of 0.15 M was used to neutralize the system. The final MD system had *∼*40,000 atoms. The residue mutations were introduced using Pymol to construct the mutant models. The wild type and mutant MD systems were subjected to minimization for 20,000 cycles. Later, the MD system was slowly heated from 0 to 300 K over a period of 3 ns in NVT and NPT ensembles. The MD systems were equilibrated for 50 ns at 300 K under NPT conditions. Finally, three independent simulations were conducted for each MD system and the final productions runs were conducted over a period of *∼*350 ns. The temperature was maintained at 300 K using the Langevin thermostat and pressure was maintained at 1 atm using the Langevin barostat. Hydrogen bonds were constrained using the SHAKE algorithm^41^ and long-range electrostatics were treated using Particle Mesh Ewald method.^42^

**Figure 2:**
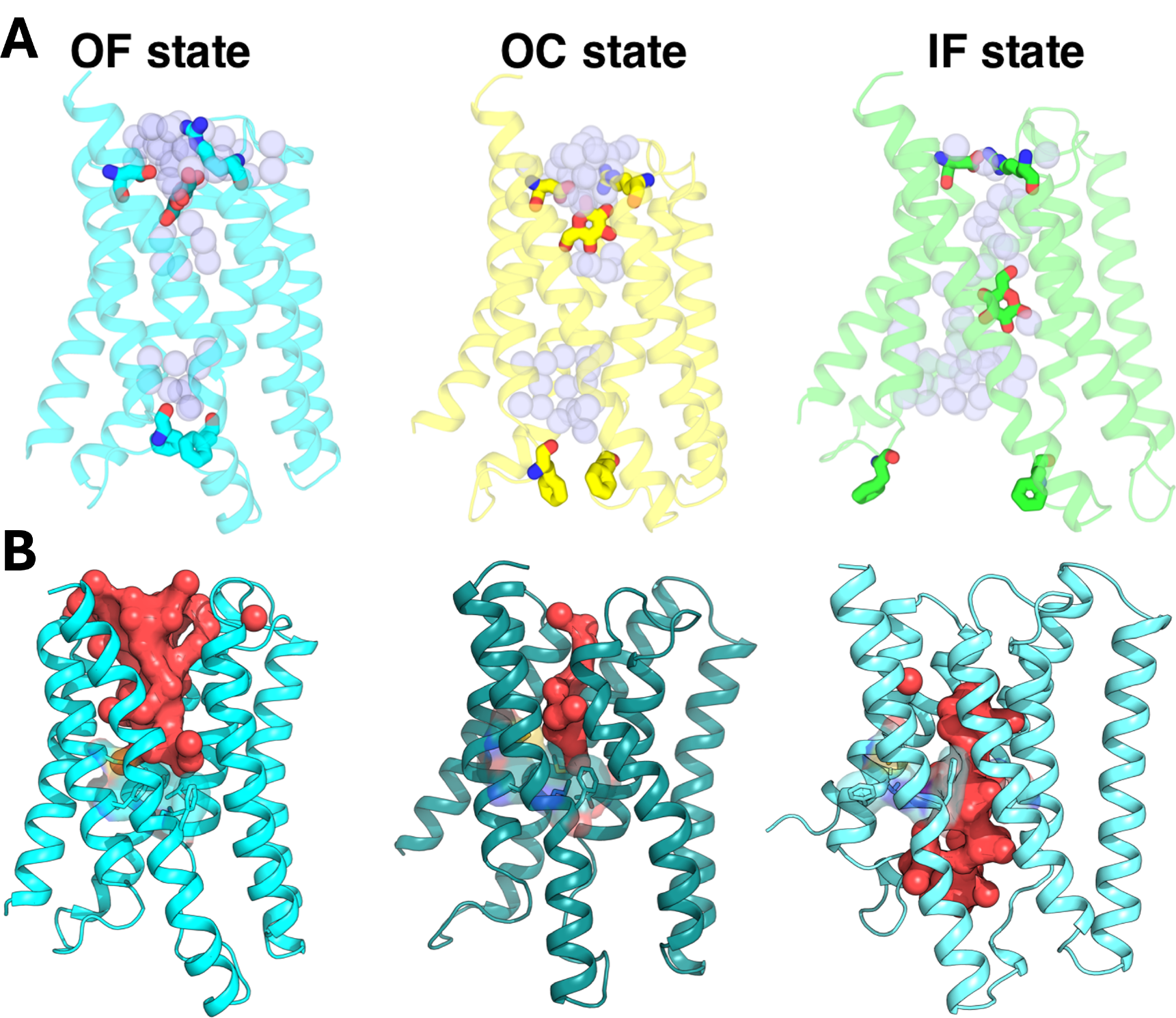
Water conducting models of holo and apo ATSWEET1. (A) Substrate bound states OF, OC, and IF of AtSWEET1. The gating residues and glucose molecule is shown in stick representation. The water molecules are shown as spheres. (B) The water filled cavities of OF, OC and IF states of apo AtSWEET1 are shown in red color, respectively. The secondary hydrophobic gating residues are represented as sticks.

### Trajectory analysis

CPPTRAJ module^43,44^ in AMBER, MDTraj^45^ and PyTraj^46^ were used for post-processing the MD trajectories. The MD snapshots were visualized and analyzed using VMD 1.9.4^47^ and Pymol v1.7.^48^ Hole program was used to calculate the pore channel radius. ^49^ The number of water molecules in the translocation pore was calculated using an in-house tcl script. The contour plots are generated by mapping the x and y coordinates of C*α* atoms of extracellular and intracellular side of the transporter.

### Yeast growth assay

AtSWEETs in the pDRf1-GW were used for yeast growth assay . Yeast growth assays were carried out as described previously with minor changes.^18,50^ The hexose transporter deficient yeast strain EBY4000 [hxt1-17D::loxP gal2D::loxP stl1D::loxP agt1D::loxP ydl247wD::loxP yjr160cD::loxP] was used for transformation with vector (negative control), Hxt5 (positive control), and AtSWEETs, respectively. For spotting assays, cells were grown in liquid synthetic medium (0.17% YNB and 0.5% (NH4)2SO4) with 2% maltose and then diluted to an OD600 of 0.1 in water. Three 10-fold serial dilutions were performed. The diluted yeasts were plated on synthetic medium containing either 2% maltose (as a control) or 2% glucose. Yeast growth was documented by Gel Doc (Biorad) after 3 days at 28 *^◦^*C.

## Results

### Molecular basis of water conduction in SWEETs

In our previous studies, we characterized the complete transport of glucose and sucrose in SWEETs and identified all functionally important states using extensive MD simulations.^20,21,25^ The observation of hydration properties in the functionally important intermediate states reveals that water molecules could rapidly access the translocation pore in the OF and IF states. The water-filled cavities of OF and OC states reveal that the water molecules are trapped at the center of the transporter and cannot enter the intracellular half (Fig. 2A, S2A and S3A). The IF state forms a continuous water conducting pathway to the intracellular side, although the extracellular side remains inaccessible (Fig. 2B and S4A). The residues Leu19, Phe20, Thr47, Met141, Phe168 and Cys172 acts as a secondary hydrophobic layer and block the water permeation in the OF state (Fig. S5). These gating residues are far away from each other in IF state and hence water molecules easily access the binding site (Fig. S5).

The binding of glucose molecules in OF state displaces roughly 20-25 water molecules to the bulk. The water displacement coupled with glucose binding is both entropically and energetically favorable. The closure of the extracellular cavity reduces the volume of the translocation pore and leads to the expulsion of most water molecules in OC state. In the glucose bound OC state, about 7-15 water molecules are identified in the pore channel. The hydrophobic residues at the center of the transporter split the transporter into two halves and thereby restrict the water permeation to the cytoplasmic side in OF and OC states. The conformational change to IF state enables the movement of the substrate to the intracellular side. The water molecules at the extracellular half coupled with glucose are co-transported to the other side of the transporter (Fig. 2A). Our study shows that *∼*7-10 water molecules are co-transported with glucose and overall cotransport of water molecules are identified as energetically favorable downhill process. The cotransport of water along with glucose confirms the validity of Hypothesis 1 in the wild-type AtSWEET1.

The salt bridge (Lys63-Asp185) at the extracellular half closes the translocation pore at the periplasmic side and restricts the motions of the helices. The water molecules transiently access the protein and permeate into the binding sites. Examination of the number of water molecules in the apo state reveals that OF state has more water molecules compared to OC and IF states (Fig. S6). The IF state has *∼*45-50 water molecules inside the translocation pore and forms a continuous pathway to the intracellular side (Fig. S6). In OC state, the water molecules are locked inside the pore channel and do not move in or out of the cavity as the transporter is closed at both the ends (Fig. S6). The average number of water molecules are lesser compared to OF and IF states. A complete transport cycle in apo state thus transports about 20-25 water molecules across the membrane, which supports Hypothesis 2. The wildtype AtSWEET1 simulations revealed that secondary hydrophobic gates restrict the entry of water molecules to access the intracellular side of transporter. The hydrophobic residues (Leu19, Phe20, Thr47, Met141 and Cys172) at the center of the transporter come close together to restrict any further diffusion of water molecules to the intracellular half of the transporter. Leu19 and Phe20 are conserved in across the SWEET family.^17^ Cys172 is asparagine in monosaccharides transport SWEETs and serine in disaccharides transport SWEETs. Similarly, Met141 is conserved among monosaccharides transport SWEETs and equivalent residue is valine in monosaccharides transport SWEETs. ^17^ These intermediate hydrophobic gates are evolutionarily conserved and observed in prokaryotic sugar transporters, SemiSWEET.^51,52^ A pair of phenylalanine residue in triple helix bundle at the intracellular side acts as a rate limiting step for the conformational transition to IF state. Previous MD studies reveal that these residues act as a pair of secondary gating residues and decrease the free energy barrier for opening the intracellular translocation pore channel. A closer examination of the simulation data reveals that water access to the other side of the membrane was restricted at the center of the transporter. In the IF state, water molecules readily diffuse to access the binding site. However, the water access to the extracellular side is not observed as the residue pairings of Lys64-Asp185, Tyr57-Asp185 and Tyr179-Asn66 form polar interactions to close the extracellular surface. Hence, we hypothesized to mutate these hydrophobic gates to a smaller residue, alanine, to investigate the water transport to the cytoplasmic side of the transporter.

We performed *in silico* single and double mutations to test whether the increase in translocation pore channel radius grants the water molecules access to the other half of the transporter. We ran three independent simulations of each mutant in OF, OC and IF states over a period of 350 ns. The mutations of hydrophobic residues to alanine increase the number of water molecules in the OF and OC state. The water-conducting snapshots in OF and OC states agree with our hypothesis 3 that the mutation of single residue Met141Ala allows the water molecules to access the intracellular half (Fig. 3A). However, the continuous water channel was still not observed (Fig. S2B and S3B). The examination of double mutant Met141Ala and Phe20Ala MD system reveals that water molecules easily diffuse to the cytoplasmic side of the transporter in OF and OC sates despite the conformational transitions to various intermediate states (Fig. 3B, S2C and S3C). The double mutation increases the number of water molecules in the cavity in OF and OC states compared to wildtype and single mutant MD systems (Fig. S7 and S8). However, the mutation has no effect on the number of water molecules in the cavity for the IF states (Fig. S4).

**Figure 3:**
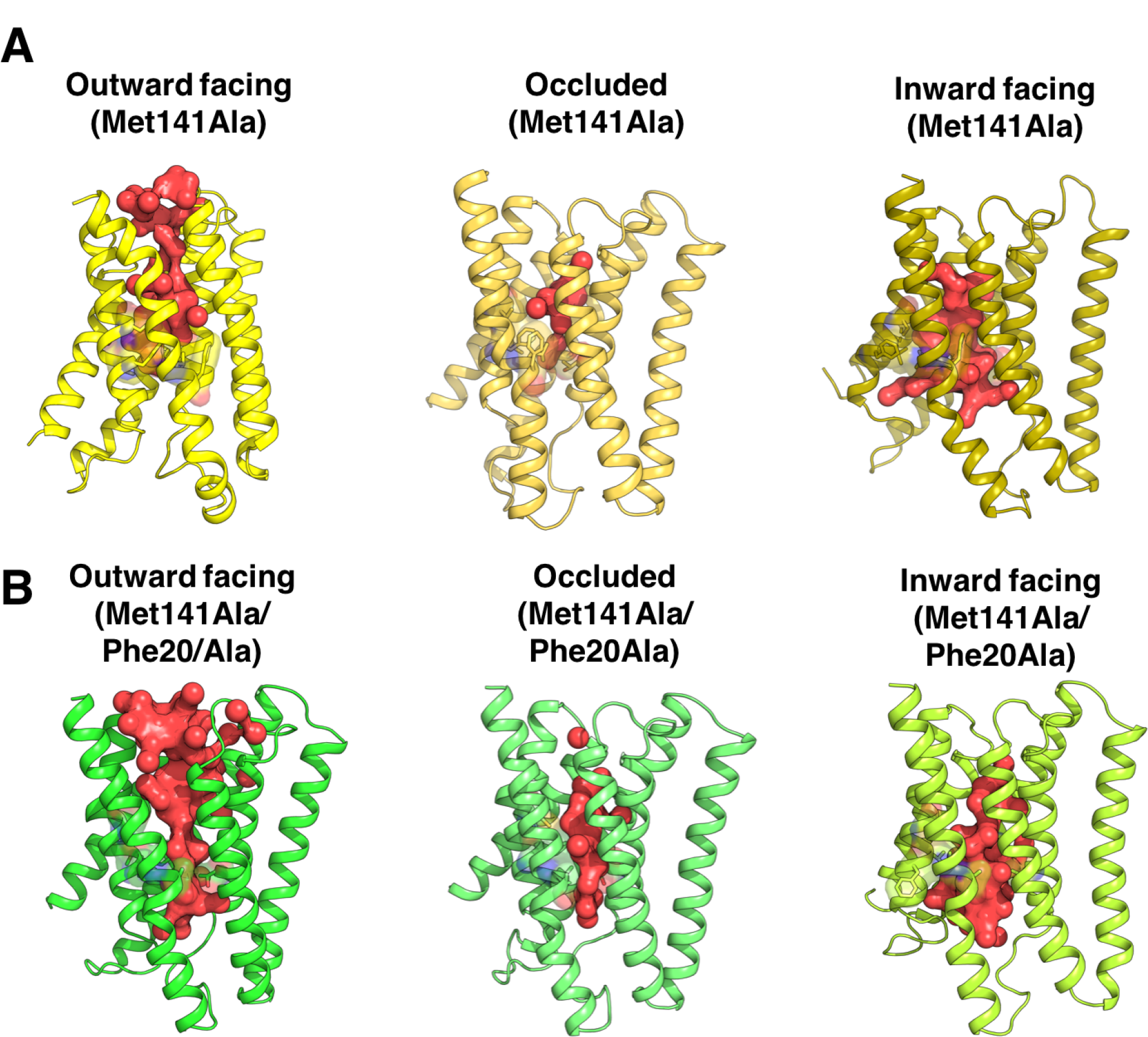
Water conducting states of Mutant systems. A) The water molecule filled cavity of Met141Ala OF, OC and IF states. B) The water permeation to the intracellular side of the transporter in the double mutant system. The water molecules are represented as red spheres.

### Substrate transport pore residues regulate water permeation in SWEETs

The water influx in SWEETs is mediated by polar and charged residues in the translocation pore. The water molecules gain access to the translocation pore by interacting with Asp65, Lys127, Arg184 and Asp185 (Fig. S9A). Further diffusion to sugar binding site was mediated by the Ser54, Tyr57 and Tyr179 residues (Fig. S9B). The residues Asn50, Asn73, Tyr142 and Asn192 in the binding site stabilize the bundle of water molecules through a hydrogen bond network (Fig. S9C). Since, the cytoplasmic cavity is closed in OF and OC states, the water molecules cannot pass beyond this site. Further, the secondary hydrophobic gates block the water permeation. In the IF state, the translocation pore is wide open and water molecules easily access the cavity; however, any exchange of water molecules from the intracellular to extracellular compartments of our membrane or vice versa was not observed in our simulation. The mutation of Met141 to alanine allows the permeation of water molecules across the hydrophobic gate. The residues Tyr142 and Thr47 mediate water entry to the intracellular side (Fig. S9D). However, the residues Pro43, Leu165, Pro145, Met161 close the pore channel along the intracellular side, thereby restricting further entry of water molecules to ultimately block the water channel in OF and OC states, respectively (Fig. S10). The double mutation Met141 and Phe20 to alanine results in the formation of a continuous water channel and allows the exchange of water molecules from extracellular to intracellular side. The formation of a water channel-like state is consistent with the hypothesis 4. The removal of hydrophobic residues allows the water molecules in the sugar binding site to pass through to the cytoplasmic side. The removal of bulky aromatic ring in Phe20 enables the interaction of water molecules with Thr26, Tyr44 and Glu80, thereby allowing diffusion to the cytoplasmic side of the transporter (Fig. 4A and 4B).

**Figure 4:**
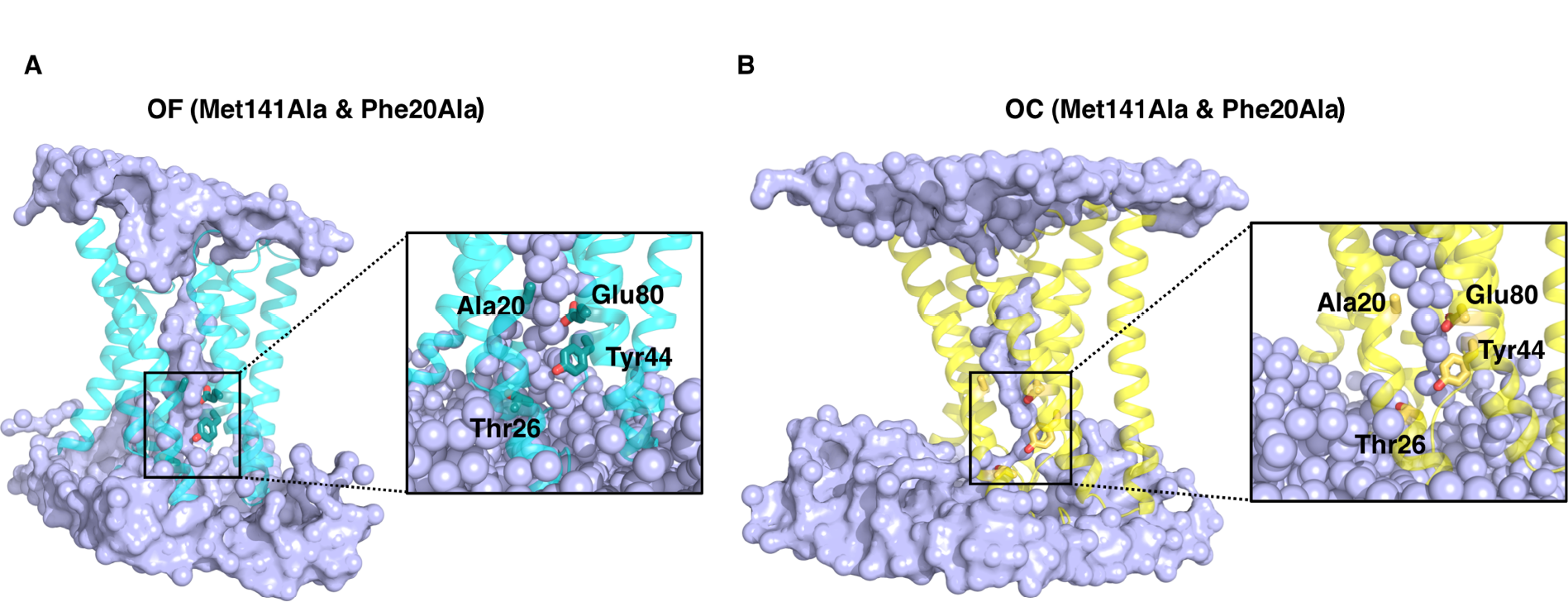
Water permeation channel. The mutation of Met141 and Phe20 to alanine leads to the continuous water flux pathway. The water conducting channel to the intracellular side via Phe20Ala was shown for A) OF and B) OC state, respectively. The water molecules are shown as blue spheres.

### Designed mutants do not affect the conformational dynamics of SWEETs

To further compare the effect of mutation on the dynamics of AtSWEET1, we analyzed the helix motions in wildtype and mutant systems. The population distribution of x, y coordinates of the C*α* atoms in each helix is shown in Figure 5. The helix dynamics plots are obtained by mapping the coordinates of the C*α* atoms of the extracellular and cytoplasmic sides onto the x-y planes. The extracellular side of the wild-type OF state reveals that helices are far apart in the transporter (Fig. 5A), whereas the inter-helix distance observed in the OC and IF states is minimized to close the pore channel (Fig. 5B-C). On the intracellular side, OF and OC conformations display a closed translocation channel while IF has an open cavity (Fig. 5A-C). Mutation of Met141 to alanine results in the movement of TM5 close to TM2 and leads to stronger helix-helix packing (Fig. 5D-E) except in IF state of the intracellular side of the transporter (Fig. 5F). The similar trend was observed in the double mutant system (Fig. 5G-I). The helix dynamics are tightly coupled between TM1 through TM4 in wildtype and single mutant systems in three different states (Fig. 5A-F). However, the dynamics are not correlated in OF and OC states of the double mutant system expect in IF state (Fig. 5G-H). This in turn shows that the removal of bulky phenyl ring in Phe20Ala facilitates the permeation of water molecules through modification of the hydrogen bond network and thus results in formation of a continuous water channel (Fig. 4A-B). In the IF state, the helices TM1-TM4 are interconnected, as observed in other MD systems and move away from TM5-TM7 to open the translocation pore (Fig. 5I).

**Figure 5:**
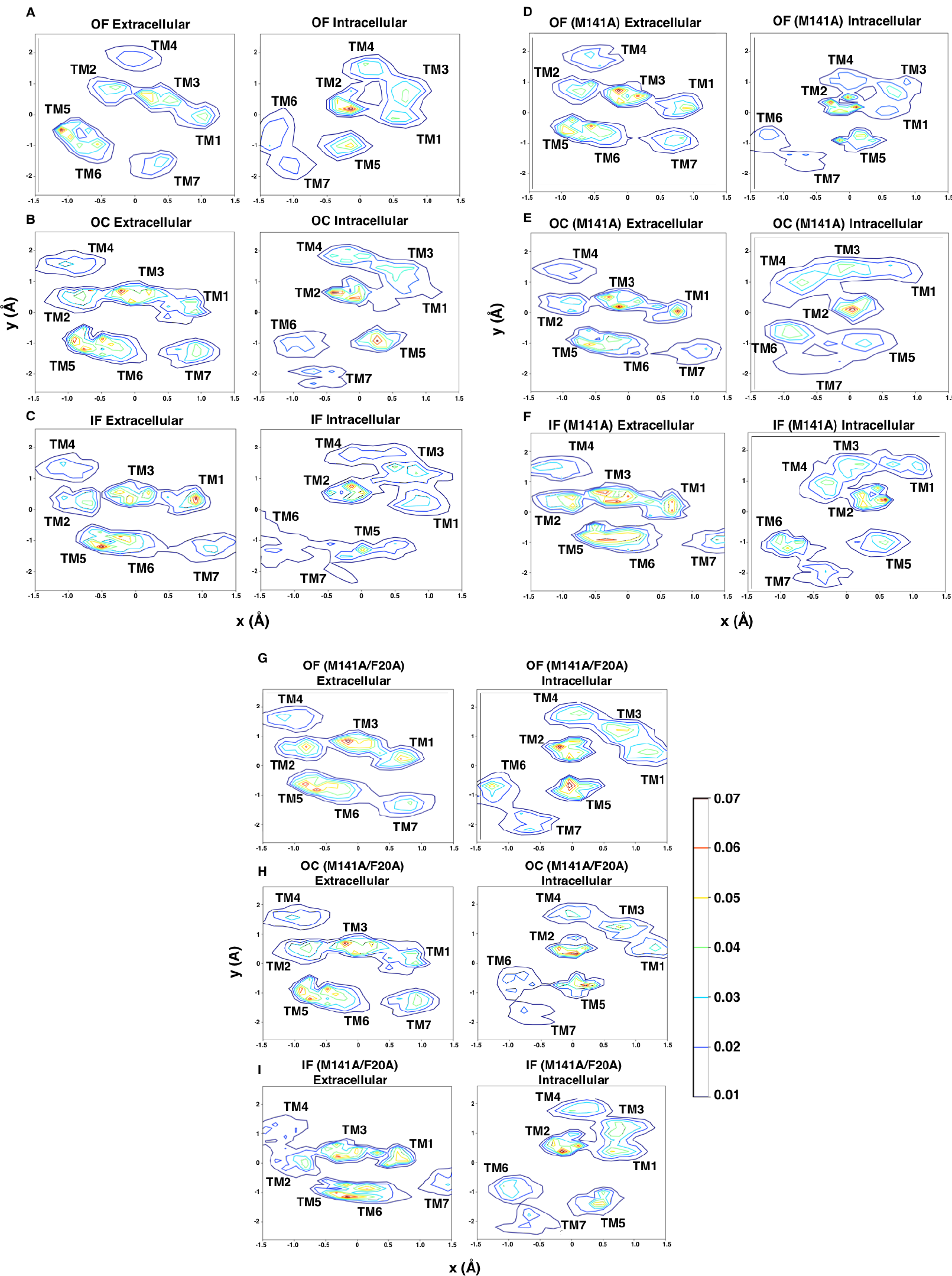
The distribution of the x, y coordinates of of the C*α* atoms for each helix in the OF, OC and IF states of wildtype (WT), single mutant (Met141Ala) and double mutant (Met141ala and Phe20Ala) of AtSWEET1. The color bar shows the probability density values for the contours.

### SWEETs transport water and H**_2_**O**_2_**

Water and H_2_O_2_ are highly related to each other and possess similar physical properties. Although the chemical behavior and van der Waals radius of H_2_O_2_ differ, the overall length and diameters of these two molecules are analogous. Since the cellular water channels, better known as aquaporins, transport both water and H_2_O_2_, we hypothesized that SWEETs may also transport H_2_O_2_. We validated our computational findings that SWEETs transport water by performing yeast growth assay using the hexose transporter Hxt5 as a control. H_2_O_2_ leads to oxidative damage to yeast cells, partially due to reduced forms of molecular oxygen, known as reactive oxygen species (ROS). Measuring water transport experimentally is challenging while measuring H_2_O_2_ transport is feasible using a yeast growth assay. The experimental results indicate that yeast with AtSWEET1, AtSWEET4, AtSWEET5, AtSWEET7, and AtSWEET8 transformed, respectively, grew normally on only maltose-containing medium. However, the presence of 0.5 mM H_2_O_2_ in the maltose-containing medium significantly affected transformed yeast cells with a SWEET except for SWEET4, which suggests that SWEETs facilitated H_2_O_2_ transport and resulted in oxidative damage and cell death (Fig. 6).

**Figure 6:**
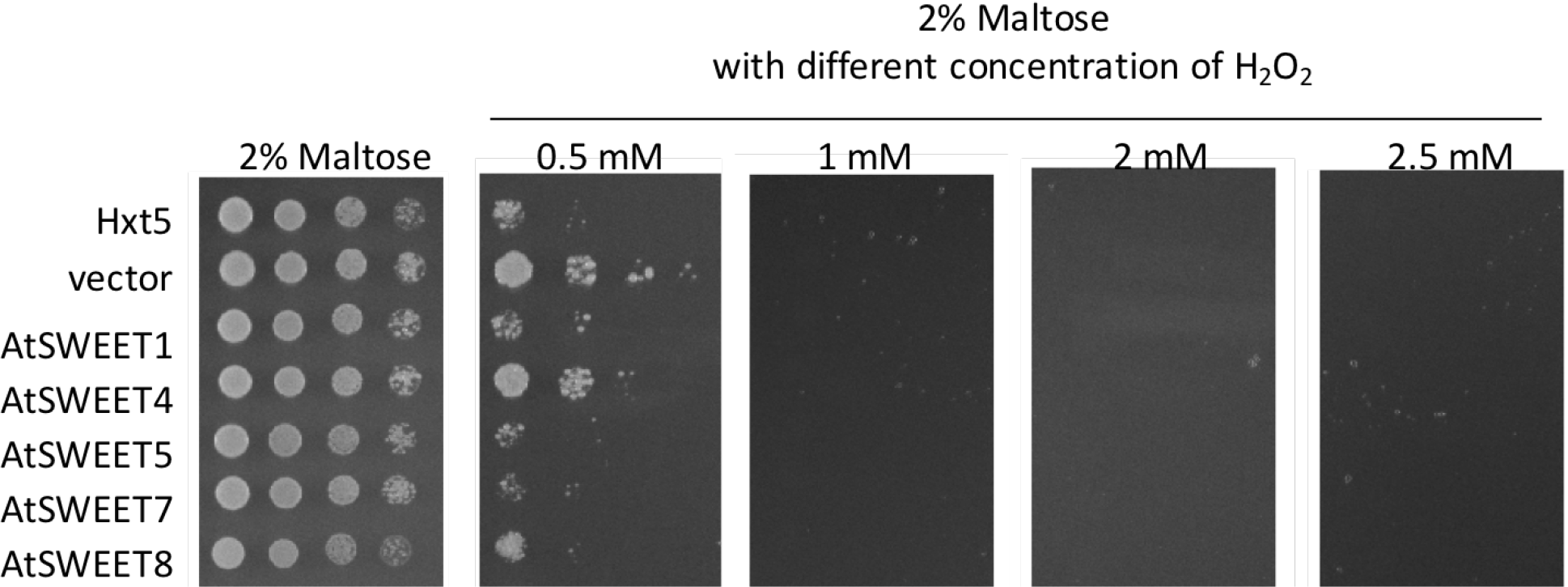
H_2_O_2_ transport by *At* SWEET family transporters estimated by complementation of the yeast growth assay.

We also characterized the transport mechanism of H_2_O_2_ in SWEET computationally by performing multiple simulations over a duration of *∼*500 ns starting from the OF state (Fig. 7A-G) to verify that H_2_O_2_ transport follows the same mechanism as water transport. Arg189 at the extracellular milieu of the transporter recognizes H_2_O_2_ and initiates the transport (Fig. 7A). The translocation pore channel is open as the gating residues are 15 °A apart and favor H_2_O_2_ binding at the periplasmic side. The coordination of H_2_O_2_ with Asp68 facilitates transport to the extra-cellular surface and forms stable contacts with Arg70 (Fig. 7B). H_2_O_2_ diffuses to the pore channel through Tyr61, Ser139, and thus initiates the closure of the transport channel at the extracellular side of the transporter. H_2_O_2_ interacts with Asn54 and Asn77 in the binding site and further entry to the intracellular site is restricted by secondary hydrophobic residues (Fig. 7C). At this juncture, the transporter is closed at both ends to obtain OC state (Fig. 7D). The increase in distance between Phe43 and Phe165 leads to the movement of helices and breakage of hydrophobic contact between secondary gating residues. As a result, H_2_O_2_ escapes from the hydrophobic site and interacts with Gln84 and Tyr44, which further facilitates transport to the cytoplasmic surface (Fig. 7E-F).

**Figure 7:**
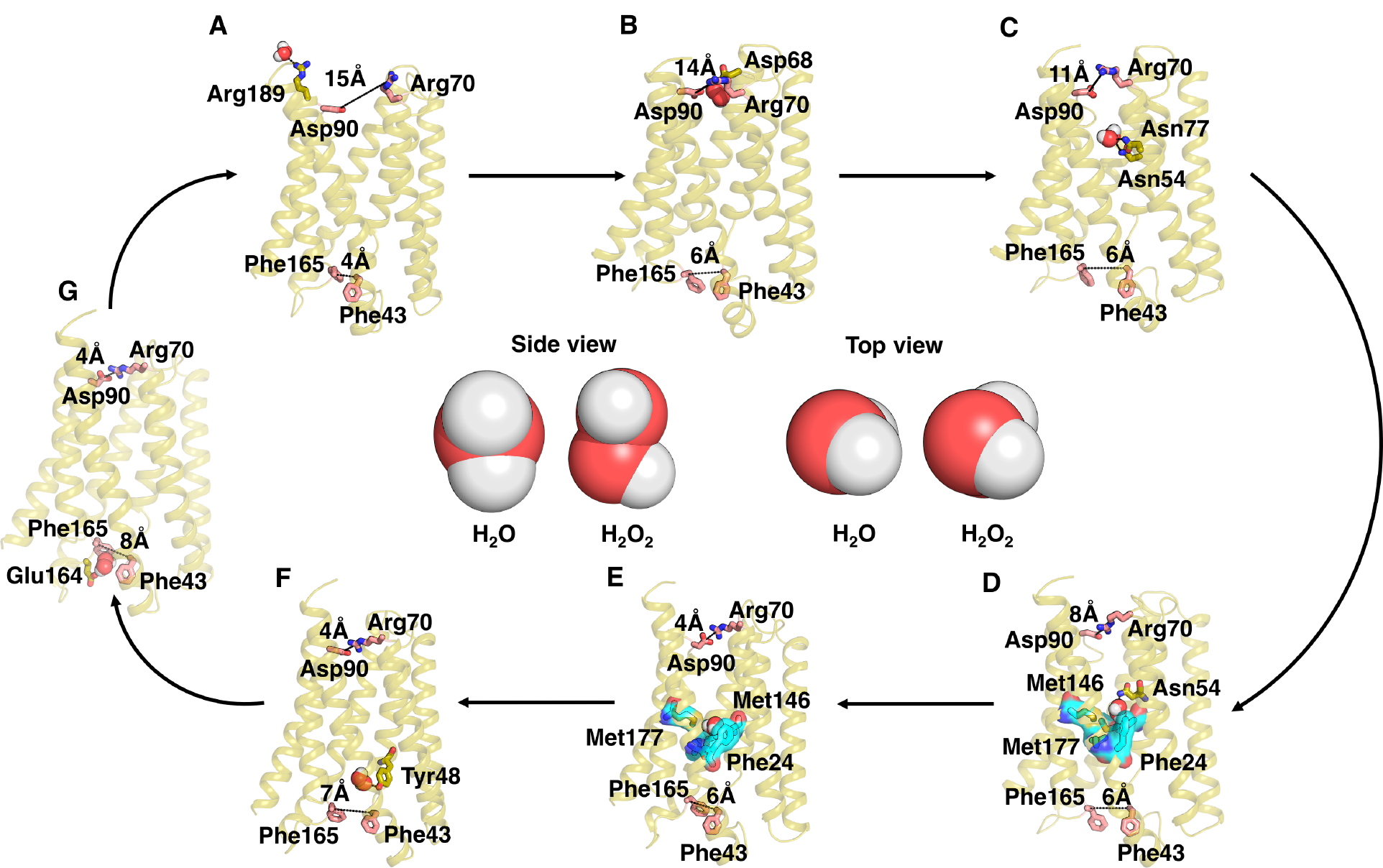
Simulation of the binding of the H_2_O_2_ molecule to SWEET. H_2_O_2_ binding mechanism and the residues that drives the transport are show in the parts A-G.

Finally, H_2_O_2_ interacts with Glu164 as the intracellular gating residue distance increases to 8 °A and leaves the transporter (Fig. 7G). Since, H_2_O_2_ is very similar to water molecules, we observed that partial opening of OC state is sufficient for the transport. The mechanism for H_2_O_2_ transport was observed to be very similar to water transport across the transporter.

## Conclusions

In this study, we identified SWEET family transporters as water co-transporters and these transporters follow different water transport mechanisms depending on the presence of substrate and mutations in the translocation pore (Fig. 1). In the substrate coupled water permeation model, water molecules are co-transported with native substrates. The examination of glucose-bound SWEET reveals that water molecules permeate to the substrate binding cavity and are coupled with the functional dynamics. Approximately 7-15 water molecules are cotransported with glucose for one complete transport cycle, proving Hypothesis 1 of substrate water cotransport model. The passive diffusion of water molecule coupled with the transport cycle is observed in apo protein thereby indicating that apo-transporter could act as a water conducting protein. The mutation of Met141 to alanine enables the free ride of water molecules to the intracellular side of the transporter. The mutation widens the water translocation channel and allows water to escape to the intracellular half of the transporter in OF and OC states as described in water escape model hypothesis. Finally, the double mutation leads to formation of continuous water channel and allows the exchange of water molecules across the membrane. The channel formed is wide enough for water to pass through without allowing permeation of the substrate, thus proving the water channel-like model hypothesis. The analysis of helical motion at the extracellular and intracellular side reveals that these mutations do not alter the conformational dynamics associated with the alternate access in AtSWEET1. Finally, the transport of signaling molecule H_2_O_2_ through SWEET1 was also observed using simulations. We performed yeast growth assays using a hexose transporter as a control and experimental studies shows that many members of the SWEET family could transport H_2_O_2_, which serves as an analogue for water transport. In summary, this study introduces a new understanding of water transport mechanisms through SWEET family membrane transporters in plants.

This study also provides avenues for further exploration of the physiological relevance, mechanism of regulation of water transport and engineering of SWEET transporters without altering their water transport function. The current study does not provide an *in planta* evidence of the impact of the water transport function on the plant physiology. Therefore, it is critical that future investigations explore the connections between drought related phenotypes and mutations that block water transport in SWEETs.^53^ Membrane transporters are often regulated via oligomerization and phosphorylation.^54–60^ In particular, SWEET family transporters form hetero- and homo-oligomers and have conserved phosphorylation sites that regulate substrate transport. ^22,53,61^ It is not clear how these regulatory mechanisms could impact the water transport function of the SWEETs. Therefore, further simulation studies are needed to establish the impact of phosphorylation and oligomerization on the water transport activity of SWEETs.^60,62^ Similarly, SWEETs have been engineered to enable selective transport of different sugars or enhance their substrate transport activity. ^63,64^ However, these engineered variants are not investigated for their water transport activity. Finally, sugar transport in plants is carried out by several different dedicated families of membrane transporters. There is a need to further investigate the water transport function of other plant sugar transporters.

## Acknowledgement

D.S. acknowledges funding from the National Institutes of Health Award No. R35GM-142745 and National Science Foundation CAREER Award NSF MCB-1845606.

## Supporting Information Available

Supplemental figures are available in the supporting information.

## Data Availability

Simulation data are available at the following web address : https://uofi.box.com/s/6jwhgyj4ehv8byvanziu9plse9aav3a2

