## Supporting Information for "SWEET family transporters act as water conducting carrier proteins in plants"

### Supplemental Figures

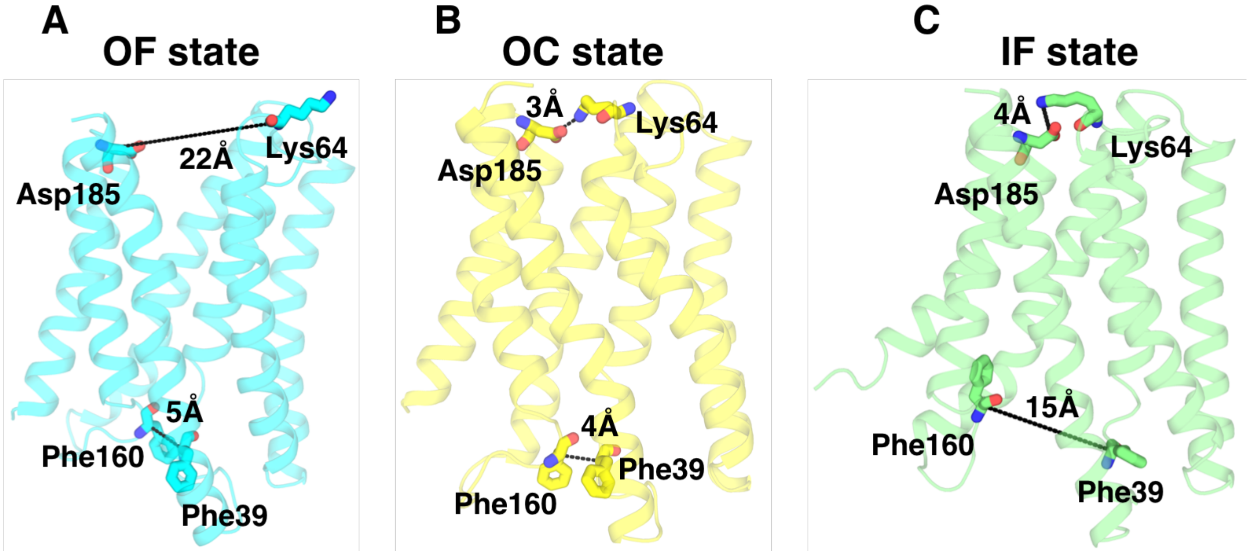


Figure 1: Homology models of ATSWEET1. The outward facing A) (OF), B) Occluded (OC) and C) inward-facing (IF) states are shown in cyan, yellow and green colors, respectively. The extracellular and intracellular gating residues are shown as sticks and the distance between them are represented as dashed lines.

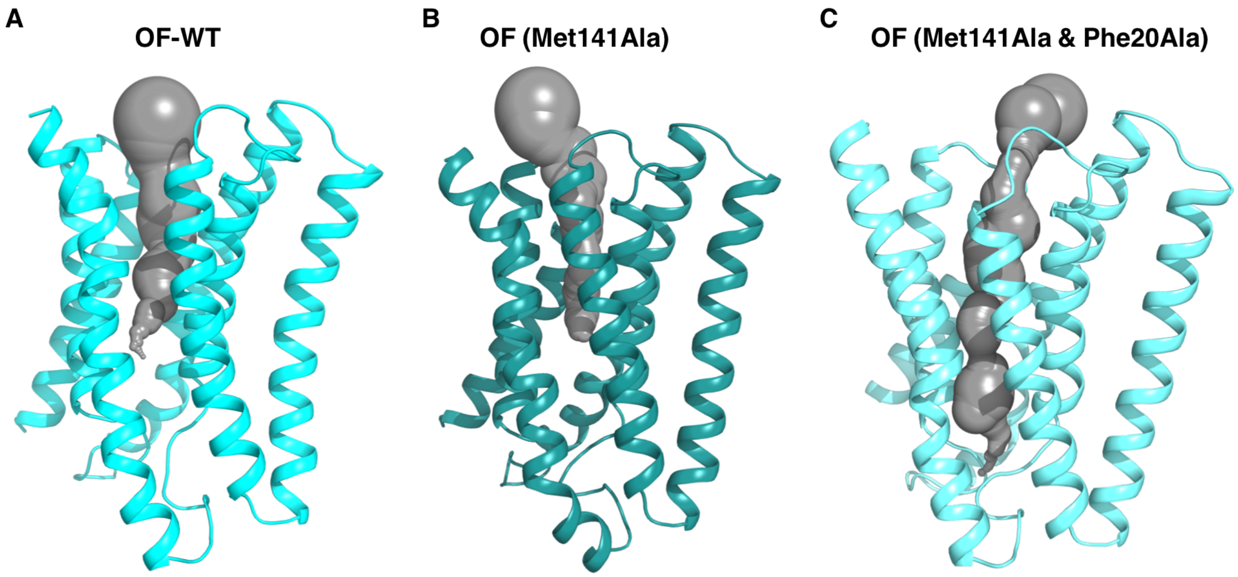


Figure 2: Translocation pore channel radius. The pore channel radius for OF state was calculated using HOLE program. The hole plots are shown for A) wildtype , B) single mutant and C) double mutant are shown, respectively.

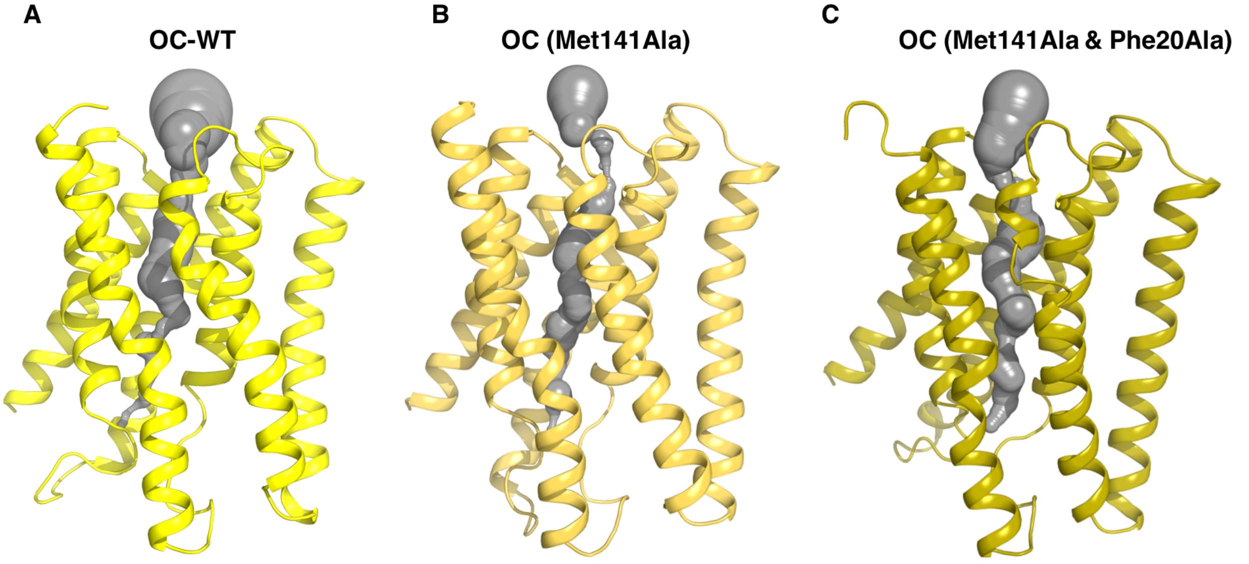


Figure 3: Translocation pore channel radius. The pore channel radius for OC state was calculated using HOLE program. The hole plots are shown for A) wildtype , B) single mutant and C) double mutant are shown, respectively.

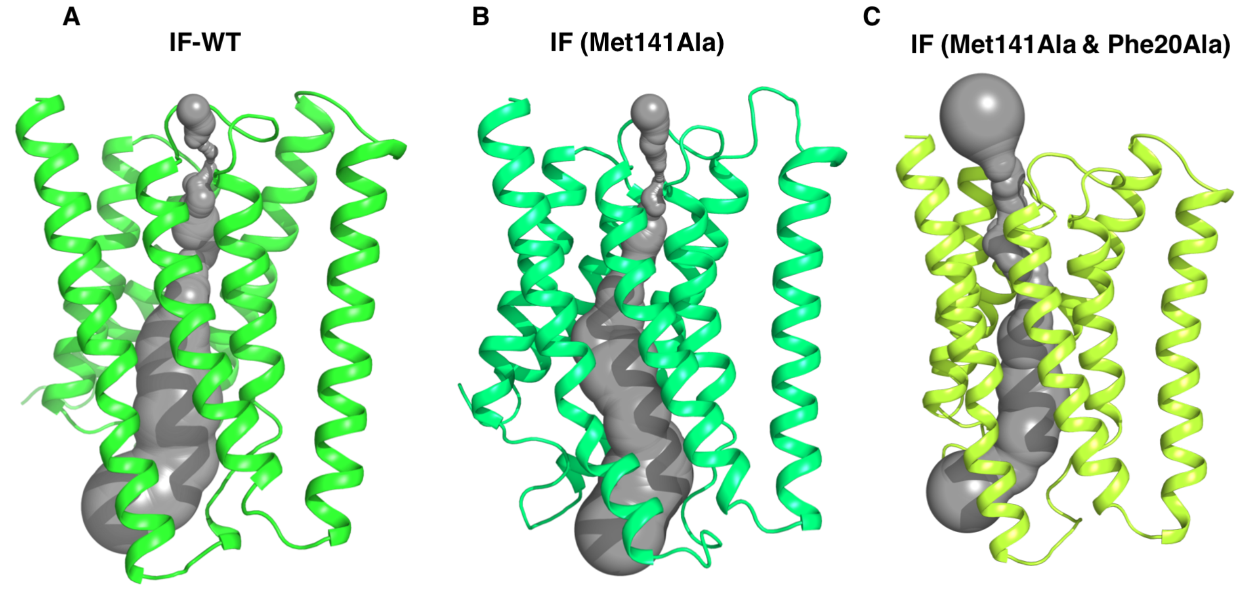


Figure 4: Translocation pore channel radius. The pore channel radius for IF state was calculated using HOLE program. The hole plots are shown for A) wildtype , B) single mutant and C) double mutant are shown, respectively.

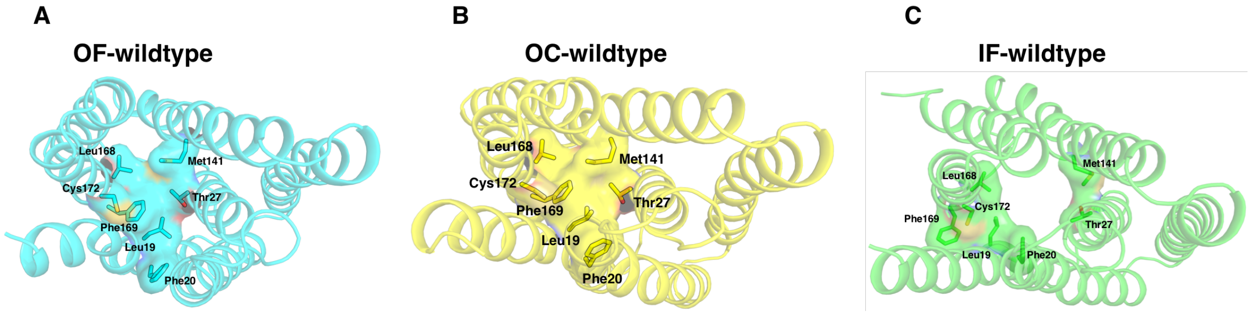


Figure 5: Secondary hydrophobic gates. The hydrophobic residues at the center of the transporter of A) OF, B) OC and C) IF states are shown, respectively.

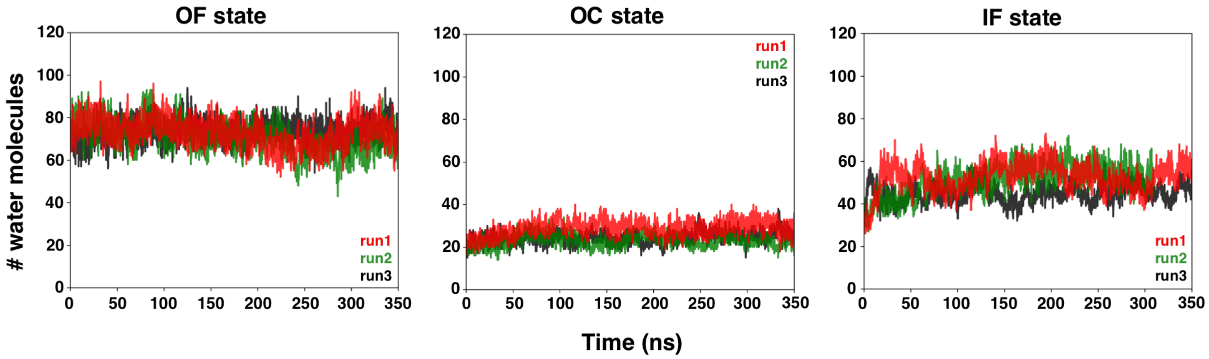


Figure 6: Water molecules in translocation pore channel of the wildtype simulation. The number of water molecules in transport pore channel in various intermediate states such as OF, OC and IF. The calculated water molecules in three different runs are shown as run1 (red), run2 (green) and run3 (black), respectively.

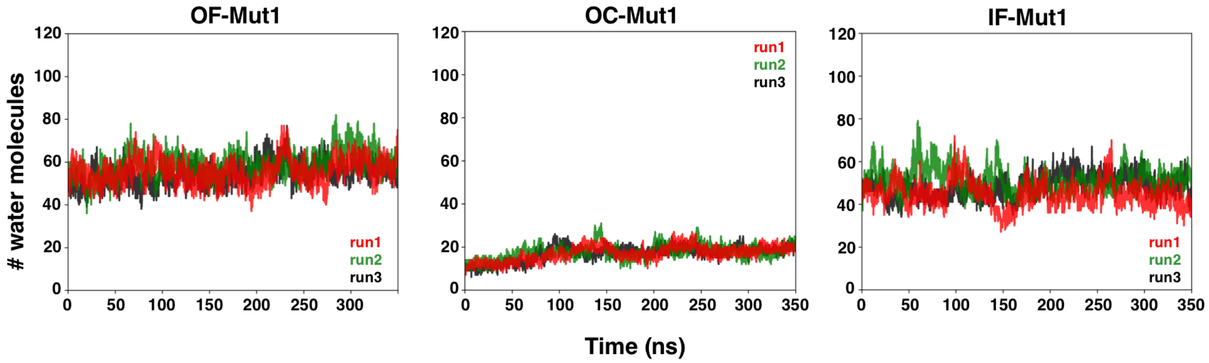


Figure 7: Water molecules in translocation pore channel of the single mutant (Met141Ala) simulation. The number of water molecules in transport pore channel in various intermediate states such as OF, OC and IF. The calculated water molecules in three different runs are shown as run1 (red), run2 (green) and run3 (black), respectively.

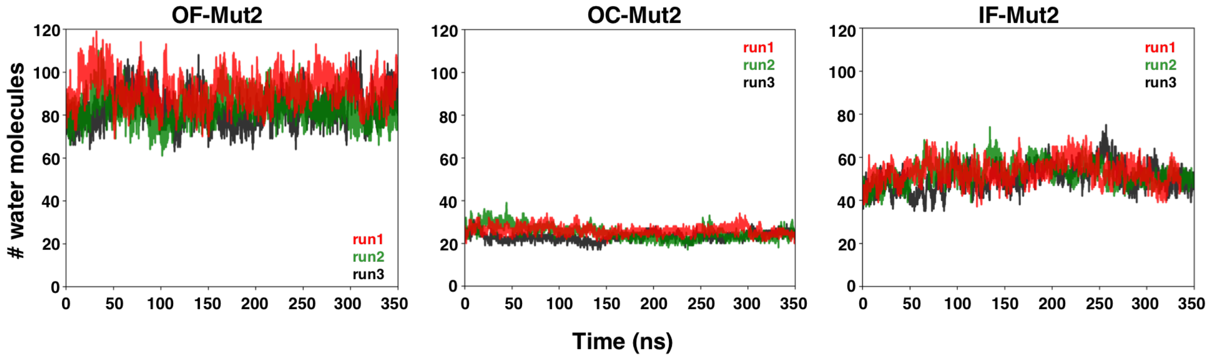


Figure 8: Water molecules in translocation pore channel of the double mutant (Met141Ala and Phe20Ala) simulation. The number of water molecules in transport pore channel in various intermediate states such as OF, OC and IF. The calculated water molecules in three different runs are shown as run1 (red), run2 (green) and run3 (black), respectively.

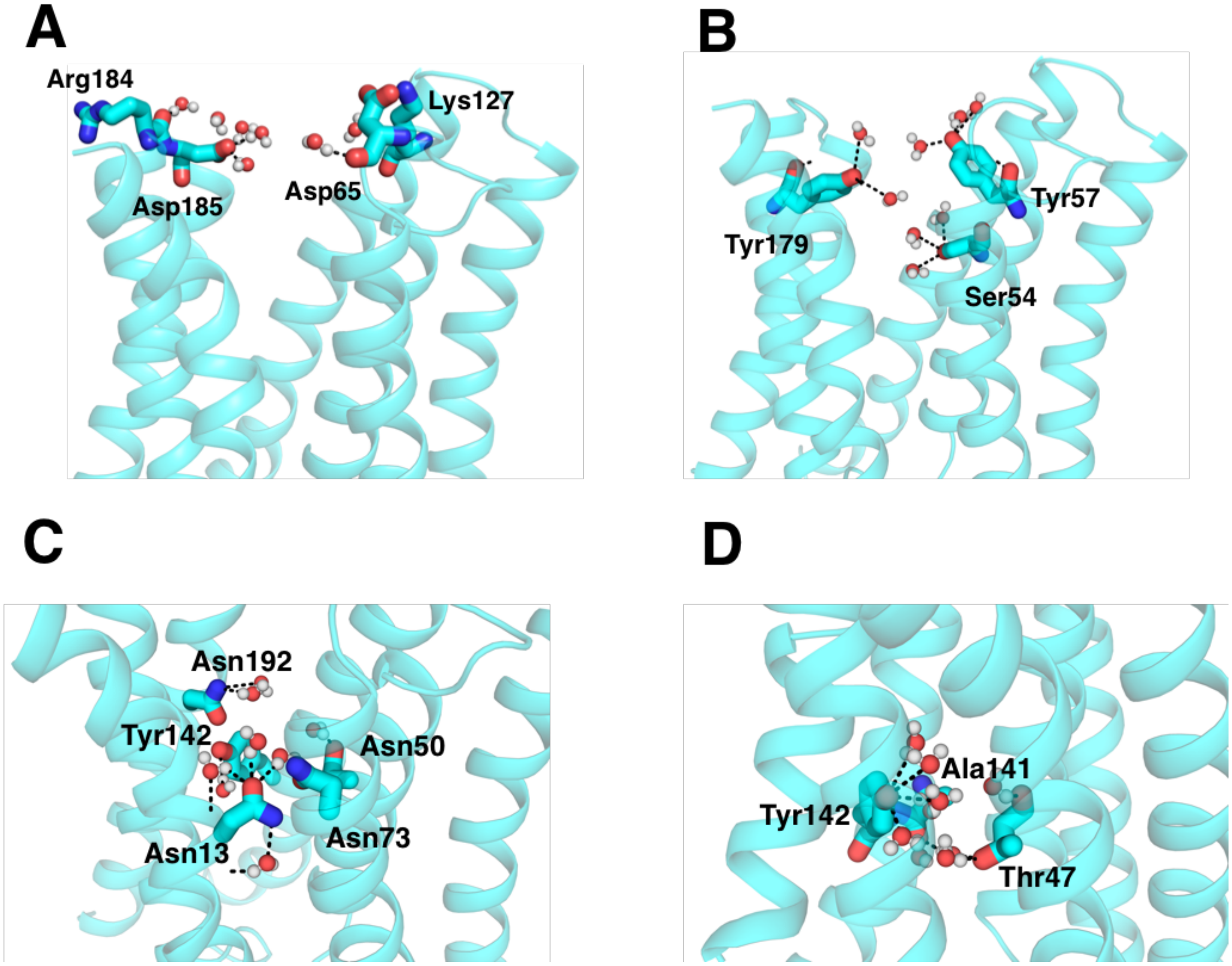


Figure 9: Residues involve in water molecule recognition, binding and translocation.

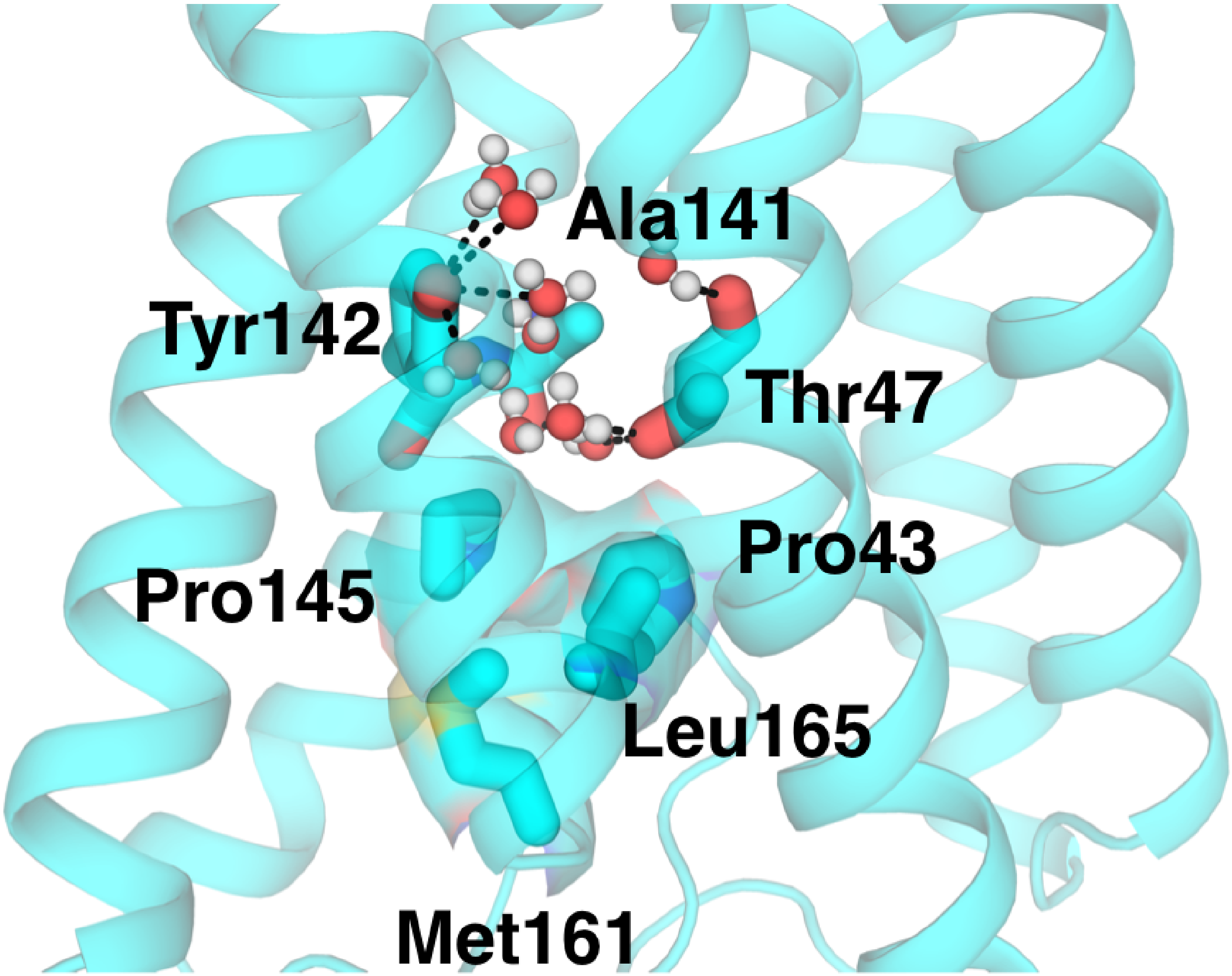


Figure 10: Mutation of Met141 to alanine allows water entry to intracellular site. Tyr142 and Thr47 forms polar contact and initiates the inward water flux. Further, entry was restricted as Pro43, Pro145, Met161 and Leu165 block the translocation pore.
